# Extensive capsule locus variation and large-scale genomic recombination within the *Klebsiella pneumoniae* clonal complex 258/11

**DOI:** 10.1101/010769

**Authors:** Kelly L. Wyres, Claire Gorrie, David J. Edwards, Heiman FL Wertheim, Li Yang Hsu, Nguyen Van Kinh, Ruth Zadoks, Stephen Baker, Kathryn E. Holt

## Abstract

*Klebsiella pneumoniae* clonal complex (CC) 258/11, comprising sequence types (STs) 258, 11 and closely related STs, is associated with dissemination of the *K. pneumoniae* carbapenemase (KPC). Hospital outbreaks of KPC CC258/11 infections have been observed globally and are very difficult to treat. As a consequence there is renewed interest in alternative infection control measures such as vaccines and phage or depolymerase treatments targeting the *K pneumoniae* polysaccharide capsule. To date, 78 immunologically distinct capsule variants have been described in *K. pneumoniae*. Previous investigations of ST258 and a small number of closely related strains suggested capsular variation was limited within this clone; only two distinct ST258 capsular synthesis (*cps*) loci have been identified, both acquired through large-scale recombination events (>50 kbp). Here we report comparative genomic analysis of the broader *K. pneumoniae* CC258/11. Our data indicate that several large-scale recombination events have shaped the genomes of CC258/11, and that definition of the complex should be broadened to include ST395 (also reported to harbour KPC). We identified 11 different *cps* loci within CC258/11, suggesting that capsular switching is actually common within the complex. We also observed several insertion sequences (IS) within the *cps* loci, and show further diversification of two loci through IS activity. These findings suggest the capsular loci of clinically important *K. pneumoniae* are under diversifying selection, which alters our understanding of the evolution of this important clone and has implications for the design of control measures targeting the capsule.

## Significance

*Klebsiella pneumoniae* is an emerging cause of multi-drug resistant hospital-acquired infections. Resistance to carbapenems, a key treatment option, is primarily associated with *K. pneumoniae* called CC258/11. Infections caused by this clone are notoriously difficult to treat. However, it may be possible to control infections by targeting the protective capsule that coats *K. pneumoniae* cells, either through capsule-specific vaccination or through treatments that break down the capsule. However, these treatments can only target a small range of capsule types. In this study we identified extensive variation in the genes responsible for capsule synthesis, indicating that *K. pneumoniae* CC258/11 change their capsule type quite frequently. This suggests treatments targeting specific capsules may be of limited use in control of CC258/11 infections.

## Introduction

*Klebsiella pneumoniae* has emerged as a common cause of multi-drug resistant (MDR) healthcare-associated infections. In particular, isolates of sequence type (ST) 258 and closely related variants, such as ST11 (as defined by multi-locus sequence typing (MLST) (1)), are distributed across continents (2-7), cause nosocomial outbreaks (3, 4, 8) and are associated with the dissemination of carbapenem resistance encoded by the *K. pneumoniae* carbapenemase (KPC) gene (2, 3, 9, 10). KPC carrying *K. pneumoniae* and other carbapenem resistant *Enterobacteriaceae* are difficult to treat (11), and have recently been recognised as an urgent public health threat by the World Health Organisation (WHO) (12), the United States Centers for Disease Control and Prevention (13), and several other government bodies.

Given the limited options for therapeutic treatment of KPC *K. pneumoniae* infections (11), there has been a resurgence of interest in *K. pneumoniae* vaccines as an alternative method of infection control, including polysaccharide vaccines targeting the capsule of outbreak strains (14). In addition, therapeutics based on capsule-targeting phage, or their capsular depolymerase enzymes, have also been proposed for control of *K. pneumoniae* (15). The polysaccharide capsule is among the most important *K. pneumoniae* virulence determinants, providing protection from phagocytosis (16), resistance to complement-mediated killing (17) and suppression of human beta-defensin expression (18). Counter-current immunoelectrophoresis techniques have distinguished 78 capsular serotypes (K types) (19, 20), however this form of serotyping is technically challenging and rarely performed (21). The genes required for capsule biosynthesis are located at the capsule polysaccharide synthesis (*cps*) locus, which shows similarity to those of *Escherichia coli* group 1 *cps* loci (22). Complete DNA sequences have been reported for only a minority of *K. pneumoniae cps* loci (23-26), however several methods of capsular typing based on genetic variation at the *cps* locus have been proposed. These include C typing, a restriction enzyme-based method that distinguishes 96 genetically distinct forms of the *cps* locus (19); and nucleotide sequencing of the conserved genes *wzi* (encoding Wzi, which anchors capsular polysaccharide to the cell surface) (27) or *wzc* (encoding Wzc, a tyrosine autokinase which polymerizes capsular polysaccharides) (28).

Recent reports have described the capsular diversity and molecular evolutionary history of ST258 (24, 25, 29-31). These studies showed that ST258 descended from an ST11-like ancestor, which acquired a 1.1 Mbp genomic region from an otherwise distantly-related ST442-like *K. pneumoniae* via recombination (24, 29, 30). The imported genomic region included a *cps* locus (*cps*_BO-4_), distinct from that present in the ST11 reference genome HS11286 (*cps*_HS11286_) (24), and a different allele of the *tonB* MLST locus (30). Compared to its ST11 ancestor, the resulting ST11-ST442 hybrid showed a change of both capsular type and ST, and has been named ST258 clade II/ST258-2 (25, 30) or ST258b (29). Subsequently, a recombination event of approximately 50 kbp arose in ST258 in which a third *cps* locus (*cps*_207-2_) was acquired by ST258-2 from an ST42-like donor, forming a new sub-lineage of ST258 named ST258 clade I/ST258-1 (30) or ST258a (29). These studies only included genomes of ST258 isolates and a small number of closely related variants (only three ST11) from a limited geographic distribution (mostly North America and Italy) (24, 25, 29, 30). Overall, just three distinct *cps* loci have been characterised in CC258/11 and two others have been indicated but not described (30).

Here we performed a genomic investigation of 39 members of the wider *K. pneumoniae* ST258/11 clonal complex (CC258/11), which were identified within a diverse collection of 230 *K. pneumoniae* genomes as well as publicly available data. Our analysis incorporates more distantly related genomes than those included in previous reports, from nine countries across four continents, including a total of nine ST11, which is the presumed ancestor of ST258. Our analysis identifies numerous large-scale recombination events within CC258/11; identifies additional STs not previously recognised as part of this clonal complex; and provides independent confirmation of the recombination events involving ST42 and ST442 in the derivation of ST258. Most importantly, we identified 11 distinct *cps* loci within CC258/11, and used these data to explore the dynamics of capsule switching and genomic variation within the clonal complex.

## Results

### Definition of the wider ST258/11 clonal complex

It is generally accepted that *K. pneumoniae* of ST258 and its single-locus variants, ST11, ST437 and ST512 represent a single clonal complex, which descended from a recent common ancestor (25, 32). We sought to identify all available genome sequences that may belong to CC258/11, therefore for the purposes of this analysis we included all draft genomes of these STs that were available in public databases at the time of investigation (**Table 1**). Additionally, MLST data and a core genome SNP phylogeny of a diverse global collection of *K. pneumoniae* (described elsewhere) (**Fig. 1**) indicated that ST395 was related to ST11: ST395 shared 4/7 MLST loci with ST11; and we estimated 0.2% genome-wide divergence between ST11 and ST395 compared to 0.6% mean divergence between ST11 and other *K. pneumoniae* KpI. We therefore included the publicly available ST395 genome 1191100241, and our own sequenced ST395 isolate K242An, in the subsequent analysis.

**Figure 1.**
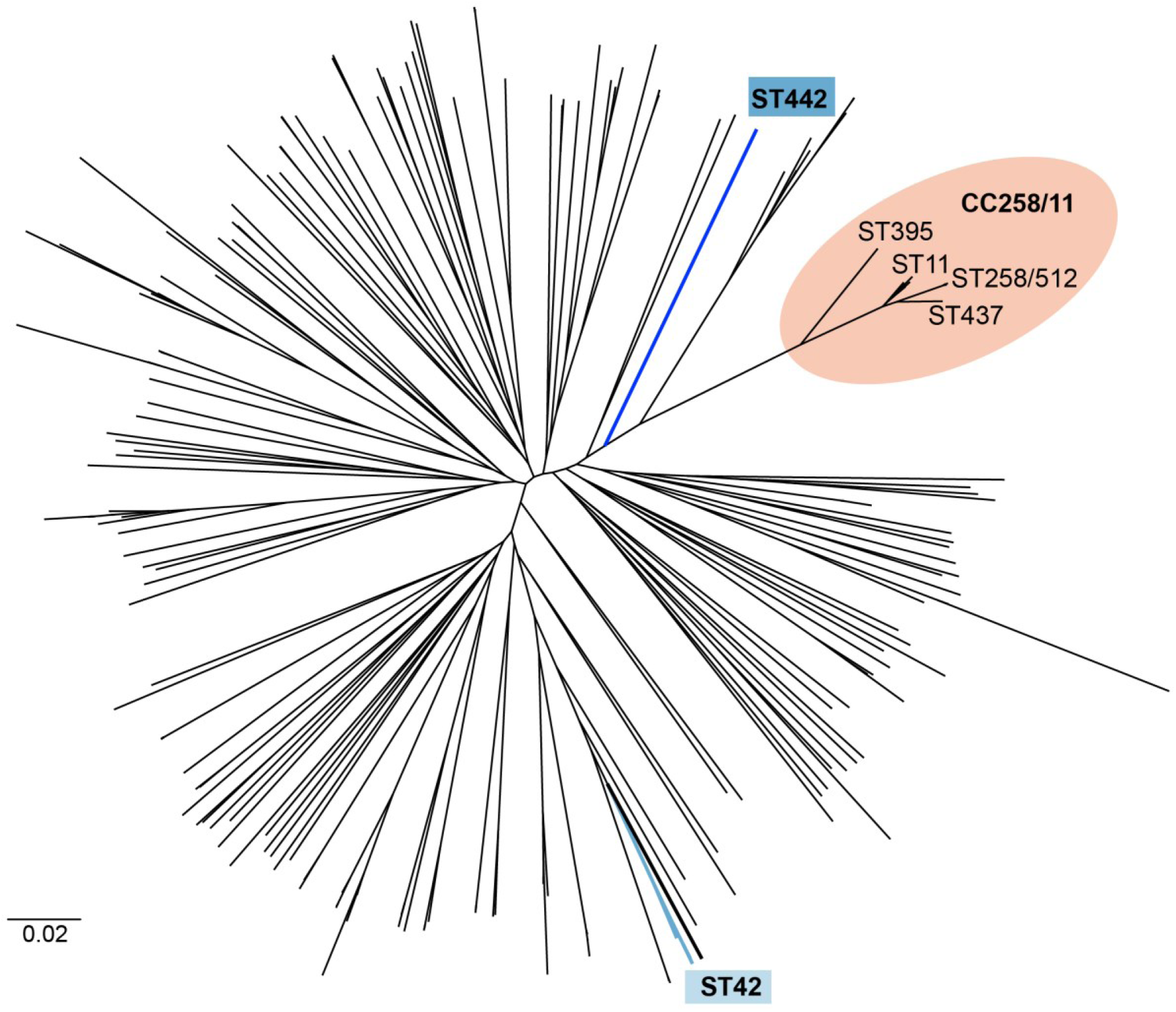
Whole-genome SNP phylogeny for 261 *K. pneumoniae* Kp I genomes. 230 genomes were from our global collection and 31 genomes were publicly available (see Methods). CC258/11 is highlighted, with sequence types (STs) marked. Genomes representing ST442 and ST42 are indicated in blue. ST442 and ST42 are the putative donors of the large-scale genomic imports resulting in the evolution of ST258-2 and ST258-1, respectively.

**Table 1.** Isolates included in BRATNextGen recombination analysis.

| Isolate | ST<sup>a</sup> | *cps* locus | K-type<sup>c</sup> | *wzi* Allele | KPC<sup>d</sup> Allele | Year | Country | Accession |
| --- | --- | --- | --- | --- | --- | --- | --- | --- |
| K242An | 395 | A | 47/NT | 105 | - | 2005 | Vietnam | ERR025564 |
| 1191100241 | 395 | A | 47/NT | 105 | - | 2011 | Netherlands | AFXH00000000.1 |
| KpOXF1 | 437 | B | 14/23* | 23* | - | 2008-2011 | UK | ERR276930 |
| KpMDU1 | 258-1 | C | NT | 29 | 2 | 2012 | Australia | AMWO00000000.1 |
| VA360 | 258-1 | Ci | NT | 29 | 2 | 2007 | USA | ANGI00000000.2 |
| ST258-490 | 258-2 | D | UNK | 154 | 3 | 2006 | Israel | ALIS00000000.1 |
| ATCC BAA-1705 | 258-2 | D | UNK | 154 | 2 | 2007 | Unknown | AOGQ00000000.1 |
| NIH<sup>b</sup> Outbreak (n=20) | 258-2 | D | UNK | 154 | 3 | 2011 | USA | AJZU00000000.1-AKAN00000000.1 |
| ST258-K26BO | 258-2 | D | UNK | 154 | 3 | Unknown | Italy | CANR00000000.1 |
| ST258-K28BO | 258-2 | D | UNK | 154 | 3 | Unknown | Italy | CANS00000000.1 |
| ST512-K30BO | 512 | D | UNK | 154 | 3 | Unknown | Italy | CAJM00000000.2 |
| HS11286 | 11 | E | 47 | 74 | 2 | 2011 | China | NC_016845.1 |
| DM23092/04 | 11 | F | 14/64/NT | 64 | - | 2004 | Singapore | ERS011902 (62) |
| DR5092/05 | 11 | G | 38 | 96 | - | 2005 | Singapore | ERS011906 |
| ATCC BAA-2146 | 11 | H | UNK | 174 | - | 2010 | USA/India<sup>e</sup> | AOCV00000000.1 |
| NCSR101 | 11 | I | UNK | 73 | - | 2007 | Vietnam | ERS011807 |
| 09-370B | 11 | J | UNK | 88 | 2 | 2009 | Vietnam | ERS012021 |
| DU10252/04 | 11 | K | UNK | 75 | - | 2004 | Singapore | ERS011907 |
| DU4033/04 | 11 | K | UNK | 75 | - | 2004 | Singapore | ERS011904 |
| DU38032/05 | 11 | Ki | UNK | 75 | - | 2005 | Singapore | ERS011911 |

<sup>a</sup> Sequence type, as defined by multi-locus sequence typing. <sup>b</sup> National Institutes of Health (USA). <sup>c</sup> As indicated by *wzi*-serotype associations reported in (27). <sup>d</sup> *Klebsiella pneumoniae* carbapenemase. <sup>e</sup> strain isolated in the USA from a patient whom had received recent medical care in India (63); NT, nontypeable; UNK, unknown serotype; \*closest allele match, where an exact match was not present in the *wzi* database.

### Evolution by recombination

We used BRATNextGen (33) to assess the extent of recombination within the CC258/11 genomes (**Fig 2**). In total 160 recombinant genomic regions were identified across the chromosome; these recombinogenic regions had a median length of 2,385 bp (range 21 bp – 1.47 Mbp; mean 57 kbp). The BRATNextGen analysis identified three large recombinant regions in the ST395 genomes, each of which was >100 kbp. The largest of these regions (which span the origin in **Fig. 2**) was approximately 1.5 Mbp in length and contained the three MLST loci that differ in ST11 (*infB*, *mdh* and *pgi*; **Fig. 2**). One of the other two recombinant regions identified in ST395 (>500 kbp in length) was shared in whole or in part by the ST11 and ST437 genomes (marked by * in **Fig. 2**). This region spans the recently reported recombination into the most recent common ancestor (MRCA) of ST258 (30); we concluded that BRATNextGen incorrectly characterised this as an import into all strains except ST258, as opposed to an import into the MRCA of ST258, simply due to a lack of resolution to differentiate these two possibilities.

**Figure 2.**
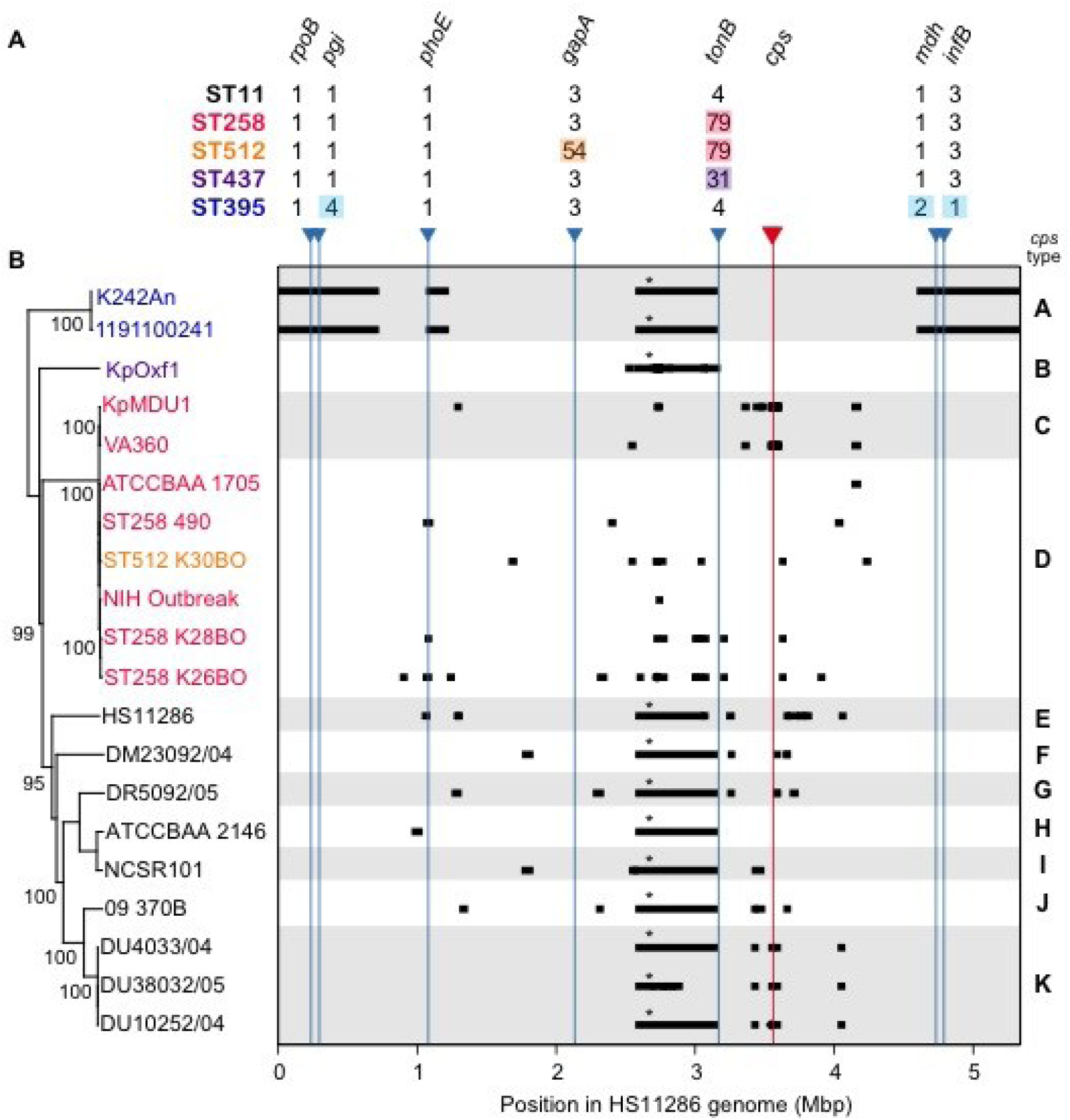
CC258/11 MLST allelic variants **(A)**, genome-wide phylogeny **(B, left)** and putative recombinant genomic regions identified by BRATNextGen **(B, right)**. Both panels share the x-axis, which indicates coordinates of the HS11286 genome (ST11) that was used as the reference for read mapping and SNP calling. Blue arrows indicate the chromosomal positions of MLST loci along this axis; red arrow indicates the position of the *cps* locus. MLST alleles for the five sequence types (STs) are shown in **(A)**; those differing from ST11 alleles are highlighted. Nucleotide polymorphisms resulting from recombinant imports are shown as black blocks in **(B)**; these were excluded from phylogenetic analysis (tree in **(B)**, bootstrap support values ≥90% are shown (%), strain names coloured as per STs in **(A)**). Asterisk indicates recombinant imports identified in whole or part in all non-ST258/512 representatives, which ChromoPainter analysis indicated was actually an import into ST258 (Fig. 3). Note that the reference genome is circular and thus what appears here as independent imports at the ends of the K242An and 1191100241 genomes, are in fact a single import spanning the origin. Grey and white background shading indicates changes in capsular locus type; types are labelled A-K on the right, genetic structures for these are given in Fig. 4.

We sought to investigate and characterise the largest putative recombination events in more detail across these isolates. We screened the collection of 230 *K. pneumoniae* genomes for genetic markers (*wzi* and/or MLST alleles) matching those within the putative imported regions. We were unable to identify any putative donors of the large ST395 recombinant regions described above, but identified candidate donors for the ST258 lineage recombination events (ST42/*wzi-29* isolate DB44834/96 and ST442/*wzi-154* isolate QMP Z4-702).

We used ChromoPainter (34) to estimate the probable ancestral origin of sites across the ST258-2 and ST258-1 genomes, as a function of our ST11, ST42 and ST442 genome sequences (**Fig. 3**). This analysis indicated ST258-2 resulted from import into ST11 of a large ST442-like sequence spanning the *cps* locus and resulting in a change of *cps* type, followed by a later import of a ST42-like 50 kbp sequence again spanning the *cps* locus, resulting in a further capsule switch and generating the ST258-1 lineage.

**Figure 3.**
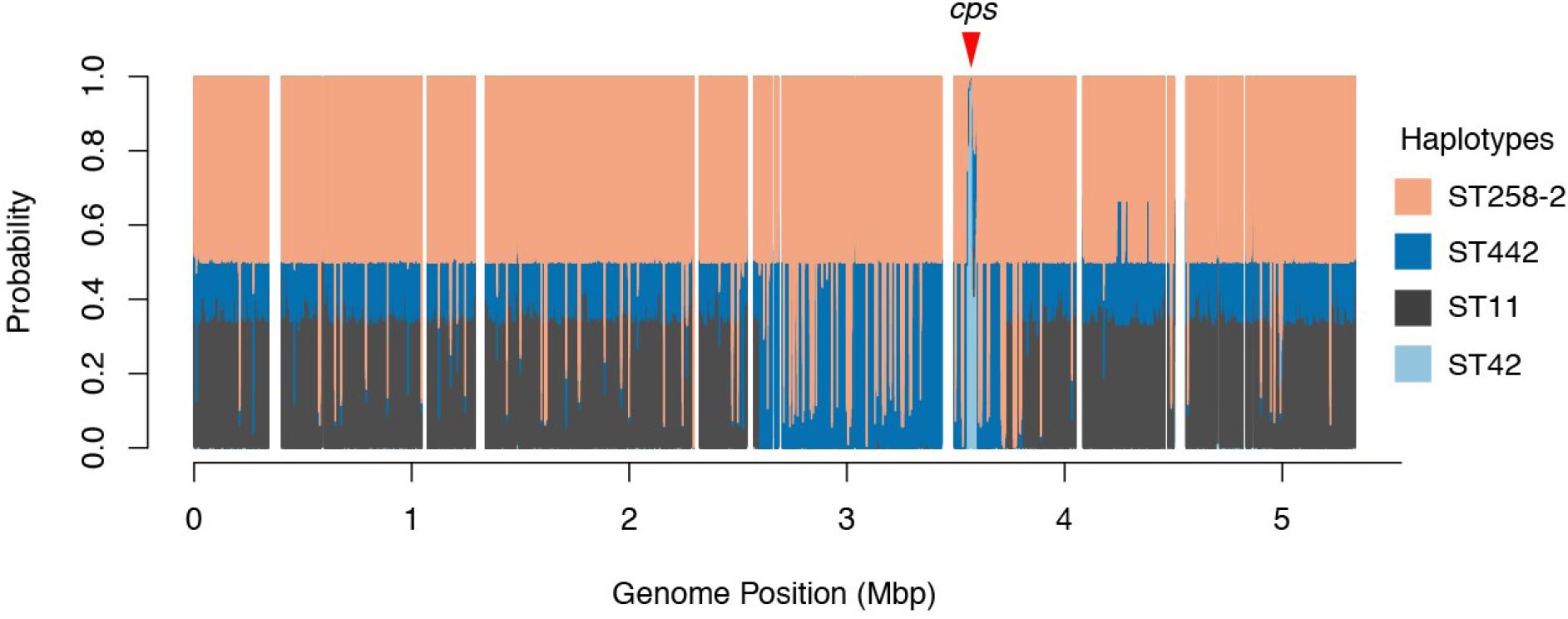
Probable ancestral origins of the ST258-1 genome, inferred using ChromoPainter to express the probability of each variable position in the genome deriving from each of ST258-2, ST11, ST42 and ST442. Most of the ST258-1 genome shares similarity with ST11 (grey) and ST258-2 (peach), with the exception of a large central segment that is derived from ST442 (dark blue) rather than ST11, and a small section spanning the *cps* (capsular synthesis) locus that differs between the two ST258 variants and is derived from ST42 (light blue). Genome coordinates as per NJST258_1 (ST258-2) reference. Note, non-variable genome positions (shown in white) were not included in analyses.

### Fine-scale phylogenetic structure

A phylogeny reflecting vertical patterns of inheritance among the wider CC258/11 is shown in **Fig. 2** (maximum likelihood tree inferred from genome-wide SNP calls after excluding SNPs introduced by putative recombination events identified by BRATNextGen). The MRCA for all ST11 genomes was close to the root of the tree, that is, the MRCA of the whole complex. Genomes of ST258 and ST512 formed a sub-cluster, within which two further sublineages were identified, which matched those previously described as ST258-1, and ST258-2 plus ST512 (25, 29). ST258/512, ST437 and ST395 were each defined by deep branches, consistent with recent independent clonal expansions of each of these sublineages. In contrast, the ST11 genomes exhibited a greater degree of diversity (**Fig. 2,** panel B). Taken together, these phylogenetic and recombination analyses indicate that members of CC258/11 descend from a common ancestor (most likely of ST11) that has diversified into several distinct lineages, some of which have novel ST combinations due to point mutations or recombination affecting the MLST loci.

### Capsular polysaccharide synthesis (cps) loci in CC258/11

We identified 11 distinct *cps* loci within CC258/11 (**Fig. 4**, **Table 1**). Three of these *cps* loci matched those previously characterised in the ST11 reference genome HS11286 (genotype *wzi-74*/*cps*_HS11286_, serotype K74) and in ST258/512 *K. pneumoniae* (genotypes *wzi-29*/cps_207-2_ and *wzi-154*/*cps*_BO-4_; unknown serotypes) (24, 25, 30). The remaining *cps* loci were annotated and are available at https://github.com/kelwyres/Kp-cps-loci.git. The capsular serotypes were predicted based on *wzi* alleles (**Table 1**).

**Figure 4.**
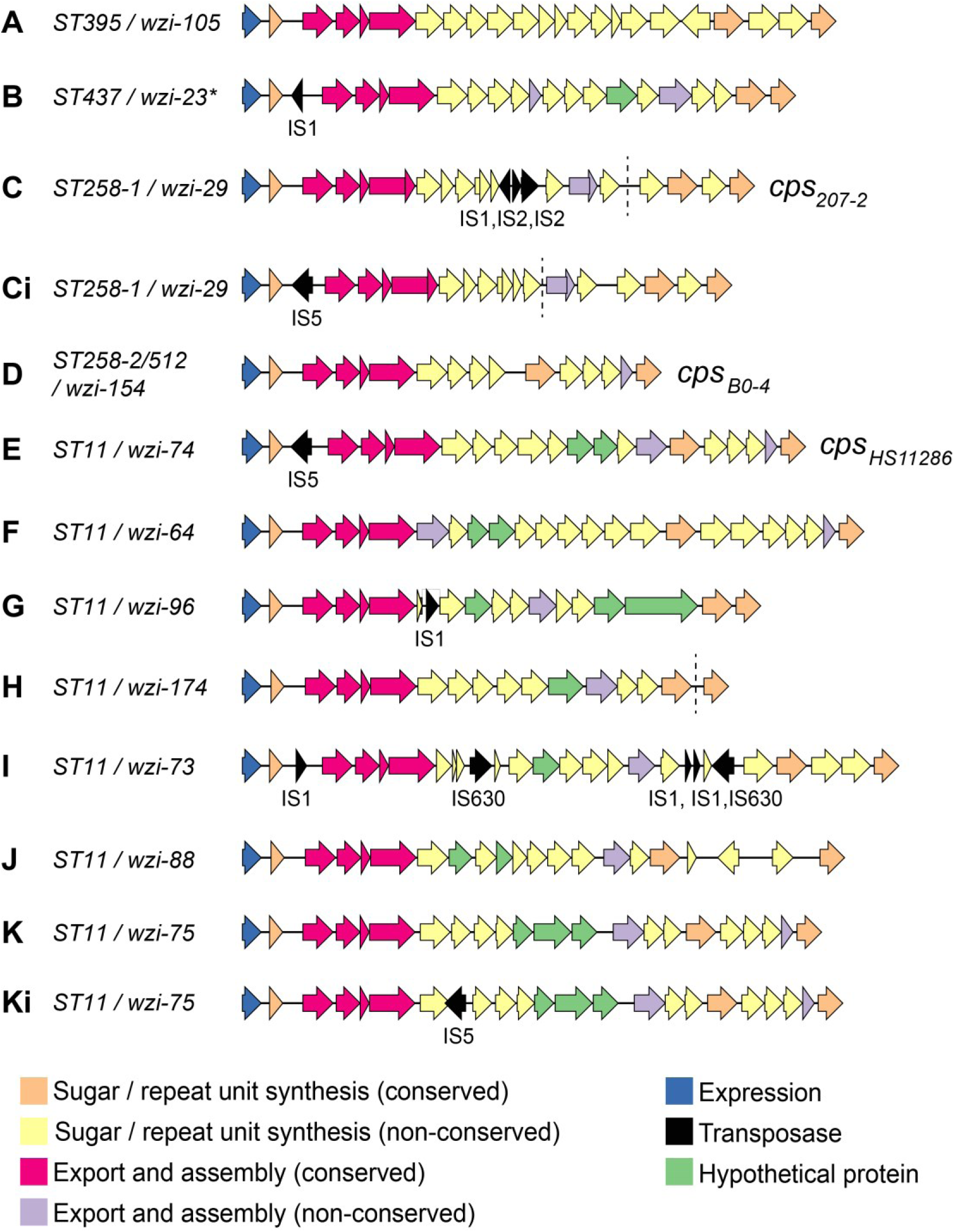
Structures of *cps* loci identified in *K. pneumoniae* CC258/11. Arrows indicate the direction, relative length and function (coloured as per legend) of protein-coding genes. Transposases were identified and labelled using the IS Finder database (https://www-is.biotoul.fr/). *Cps* loci are labelled A through K as referred to in the text, Table 1 and Figure 2; STs and *wzi* alleles are indicated (* indicates inexact match to previously defined *wzi* alleles); loci previously named in (24) are indicated (*cps_B0-4_, cps_207-2_* and *cps_HS11286_*). *Cps* loci that vary from another only in the content and/or position of transposases are indicated by ‘i’. Annotated sequences are available at https://github.com/kelwyres/Kp-cps-loci.git. Dashed lines indicate contig breaks in assemblies of three loci.

All CC258/11 *cps* loci shared a conserved macro-structure consistent with that previously reported among *K. pneumoniae* (23) (**Fig. 4 and 5**). The macro-structure was comprised of eight conserved protein-coding sequences (CDSs) situated at either end of the *cps* locus: *galF* (UDP-glucose pyrophosphorylase), *orf2* (putative acid phosphatase), *wzi, wza, wzb* and *wzc* (polysaccharide polymerisation and export) at the 5’ end, and *gnd* (6-phosphogluconate dehydrogenase) and *ugd* (UDP-glucose 6-dehydrogenase) at the 3’ end. The median pairwise nucleotide similarities within these genes ranged from 99% (*galF*) to 55% (*wzc*), with greatest genetic conservation observed at the terminal ends of the locus (**Fig. 5**). The conserved CDSs located at either end of the locus had a G+C content of >50%), similar to the rest of the *K. pneumoniae* chromosome (the overall G+C content of the HS11286 chromosome was 57.5%). In contrast, the non-conserved CDSs in the centre of the *cps* loci had <50% G+C content (**Fig. 5**). These data suggest that the evolutionary origins of the non-conserved CDSs are distinct from those of the conserved CDSs and the rest of the *K. pneumoniae* chromosome. Presumably the central CDSs have been transferred horizontally into the centre of the locus and then exchanged through homologous recombination mediated by sequence conservation in the outer CDSs.

**Figure 5.**
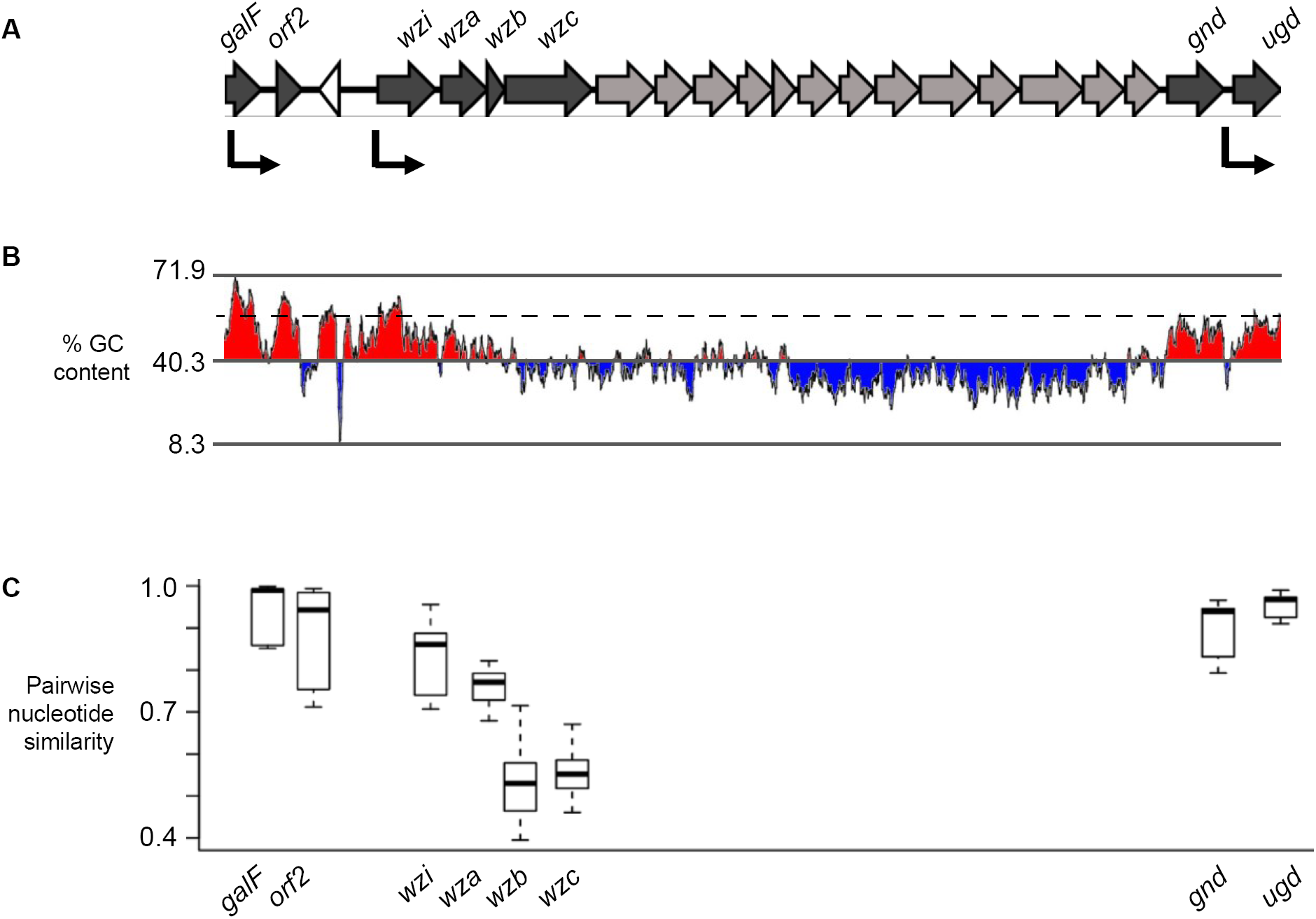
Macro-structure and features of *K. pneumoniae cps* loci. **(A)** Structure of a representative *cps* locus, *cps*-B (individual structures *cps* A-K are given in Fig. 4). Filled arrows indicate size and direction of protein-coding genes; dark grey: conserved CDSs present in all *cps* loci (gene symbols labelled); light grey: non-conserved CDSs (present in *cps*-B but not other *cps* loci); white arrow: transposase. Black arrows indicate the relative start position and direction of transcriptional units as described in (23). **(B)** G+C content across *cps*-B; red and blue indicate above and below the mean G+C content of the locus. Dashed line indicates mean G+C across the whole genome (57%). **(C)** Distributions of pairwise nucleotide similarities amongst CC258/11 *cps* loci, for conserved genes only.

The *cps* sequences indicated that variation in the central region drives differences in the overall length and CDS content of the *cps* locus, which ranged from 20 to 31 Kbp and from 16 to 24 CDSs, respectively (excluding transposases, see **Fig. 4**). Among the 223 independent CDSs annotated across the 11 CC258/11 *cps* loci, predicted protein products included sugar production/processing proteins (n = 69), sugar transferase proteins (n = 69), capsule export and assembly proteins (n = 44), sugar transport proteins (n = 15), unknown proteins associated with capsule production (n = 11) and hypothetical proteins (n = 15) (see **Fig. 4**). The non-conserved CDSs in the centre of the locus were predominantly associated with capsule-specific sugar synthesis and assembly. For example, *cps*-A, -F and -I carried *manB* and *manC* that encode a phosphomannomutase and a mannose-1-phosphate guanylyltransferase, respectively. In total, 134 CDSs were identified within the non-conserved *cps* regions. BLAST comparisons to nucleotide and protein sequences in the NCBI database indicated 58 of these proteins were associated with the synthesis, processing and/or export of specific sugars: mannose (n = 21), galactose (n = 9), glucose (n = 9), colonic acid (n = 8), pyruvic acid (n = 5), acetic acid (n = 4), fucose (n =1) and hyaluronic acid (n = 1). A further 61 CDSs had ≥70% amino acid identity to sequences commonly associated with sugar synthesis, processing and/or export. Fifteen CDSs did not match any known nucleotide or protein sequences in the NCBI database or matched sequences annotated as hypothetical. Notably, the non-conserved region of the *cps* locus of NIH outbreak-associated isolates included two putative rhamnosyltransferase genes, consistent with the reported detection of rhamnose derivatives in the capsule of outbreak-associated isolates (35).

Thirteen transposase-associated CDSs were additionally identified within the *cps* sequences (**Fig. 4**). These included IS1 (present in four *cps*), IS2 (present in one *cps*, adjacent to an IS1 insertion), IS5 (present in three *cps*) and IS630 (present in one *cps*). Furthermore, four of the IS insertions (two IS1, two IS5) occurred upstream of *wzi*, near the transcriptional start site for the majority of capsular synthesis genes. These transposases have strong promoters and are in frame with the *cps* CDS. The other IS insertions all occur within the central sugar processing regions; two of these disrupt CDS (IS1 in *cps*-G and IS5 in *cps*-Ki).

For two *cps* loci, transposase insertions were differentially present, generating variant forms of the *cps* locus. *Cps*-K (*wzi*-75), identified in three closely related ST11 isolates from Singapore, carried an IS5 insertion in the sugar-processing operon in one isolate. Both available ST258-1 references carried a copy of the *cps*-C (*cps*207-2) locus with either an IS5 insertion upstream of *wzi* or an IS1 and two IS2 insertions within the central sugar-processing region. The *cps*-C locus in the donor strain ST42 and all ST258-1 strains reported in (25) carried the IS1/IS2 insertion without the IS5 insertion. It is likely that the *cps*-C sequence imported into ST258 from ST42 was in the form of *cps*-C (containing the IS1/IS2 insertion), but has since diversified though loss of the IS1/IS2 insertions and acquisition of the IS5 insertion in some strains (*cps*-Ci).

### Phylogenetic distribution of cps loci and capsule switching

The various *cps* loci were confined to distinct phylogenetic sub-clusters within CC258/11 (**Fig. 2**). Across the clonal complex, genomes with different *cps* regions were also differentiated in terms of nucleotide divergence across conserved regions of the genome (**Fig. 6**). As presented above, the ST258/512 cluster harboured two *cps* loci, each confined to one of the ST258 sub-lineages that were separated by a few hundred SNPs. The next closest genome pair with different *cps* loci was NCSR101 and ATCC BAA-2146 (both ST11), which differed by 500 SNPs across the rest of the genome, whereas all other pairs of genomes with different *cps* loci differed by >1,000 SNPs. These data suggest that stable capsule switching events may occur as frequently as one in every 102-103 nucleotide substitutions in *K. pneumoniae*. Extensive differences in gene content were also observed within CC258/11 and were correlated with nucleotide divergence (**Fig**. **6**). Genomes with different *cps* loci also differed substantially in terms of gene content outside the *cps* locus (mean 809 genes different between pairs of genomes).

**Figure 6.**
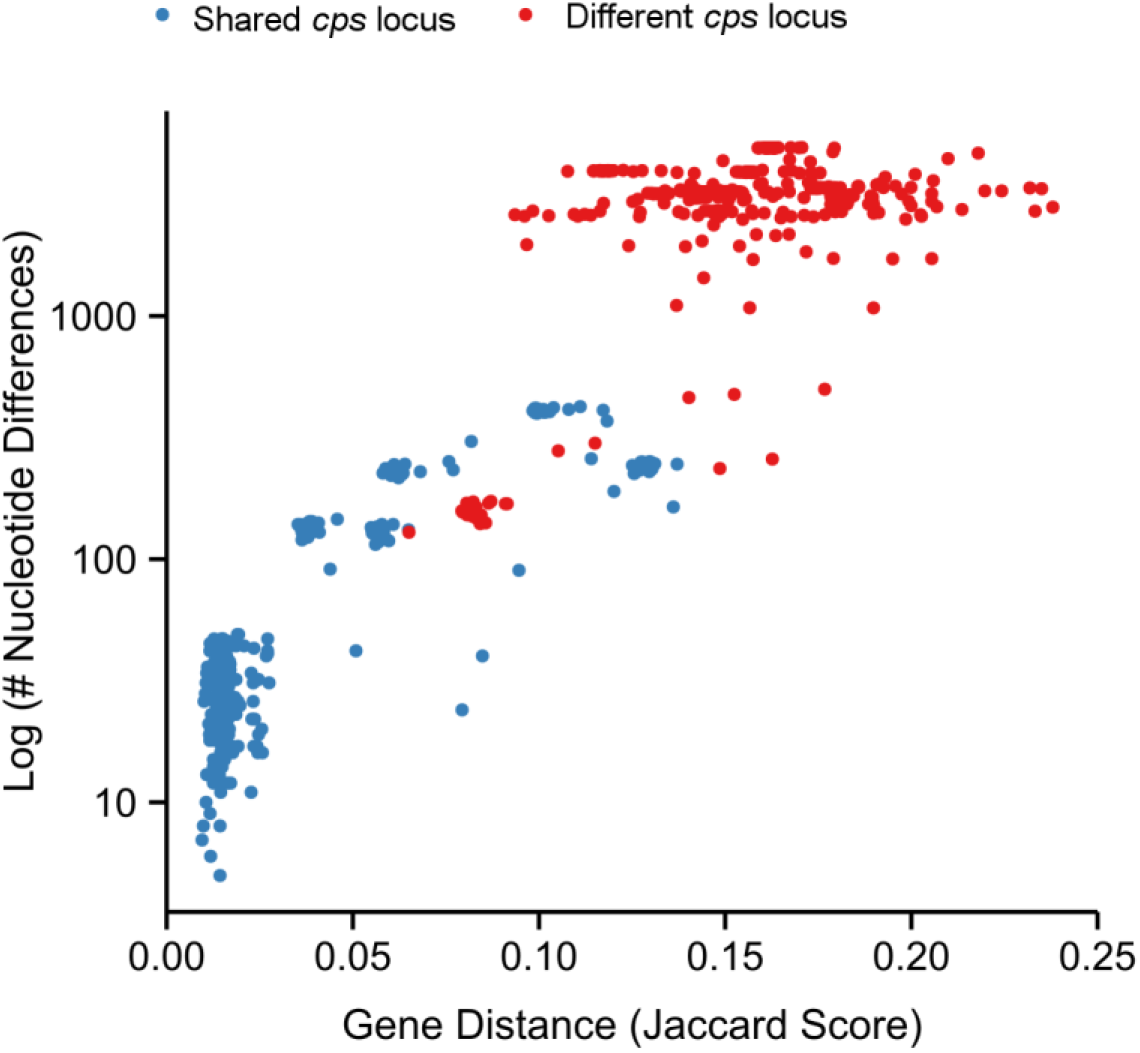
CC258/11 Total nucleotide differences vs. gene content distances (Jaccard scores) for pairs of CC258/11 *K. pneumoniae*. Points are coloured blue for genome pairs that share a *cps* locus structure, red otherwise.

## Discussion

Our data provide important insights into the evolution and capsular diversity of globally distributed KPC-associated *K. pneumoniae* CC258/11. Our analyses support previous evidence that ST258/512 emerged as a hybrid lineage within this clonal complex (29, 30) and, through use of an alternative analytical framework (ChromoPainter) and novel genomes, we provide independent confirmation of the large-scale recombination events that drove its emergence and resulted in capsule switching. Significantly, we show that these large-scale recombination events and capsule switches are not unique events in the history of this clonal complex, but are a major and potentially common driver of variation within CC258/11.

The *K. pneumoniae* ST442/*wzi-*154 strain in this study was isolated from a case of bovine mastitis at a dairy farm in New York State in 2006 (36), which is geographically close to the location of the first reports of ST258 in New York in 2000 (37). The ST42/*wzi-*29 was isolated in 1996 from a blood stream infection in a hospital in Singapore (38). This evidence supports the notion that ST42/wzi-29 was present in South East Asia, where ST11 is most common.

We identified an additional large (1.5 Mbp) recombination import and a second import of approximately 113 kbp within the ST395 genomes (**Fig. 7**), which lay within CC258/11 (**Fig. 2**). These large-scale recombination events led to the acquisition of divergent alleles at three of the seven *K. pneumoniae* MLST loci (**Fig. 7**). As a consequence, ST395 (a 4-locus variant of ST258) has not routinely been considered part of the epidemic CC258/11, even when KPC ST395 were found co-circulating with KPC ST11 in Asia (10). Our data indicate ST395 is highly similar to ST258 in that it is a hybrid KPC-associated strain emerging within CC258/11. Our analysis therefore provides strong evidence for an expanded definition of CC258/11 to include all strains that share a MRCA with ST11 as opposed to being based on shared MLST alleles (**Fig. 2**). Furthermore, our data suggest that all CC258/11 isolates, including ST395 and ST437, should be included in any surveillance and research investigations focussed on KPC CC258/11.

**Figure 7.**
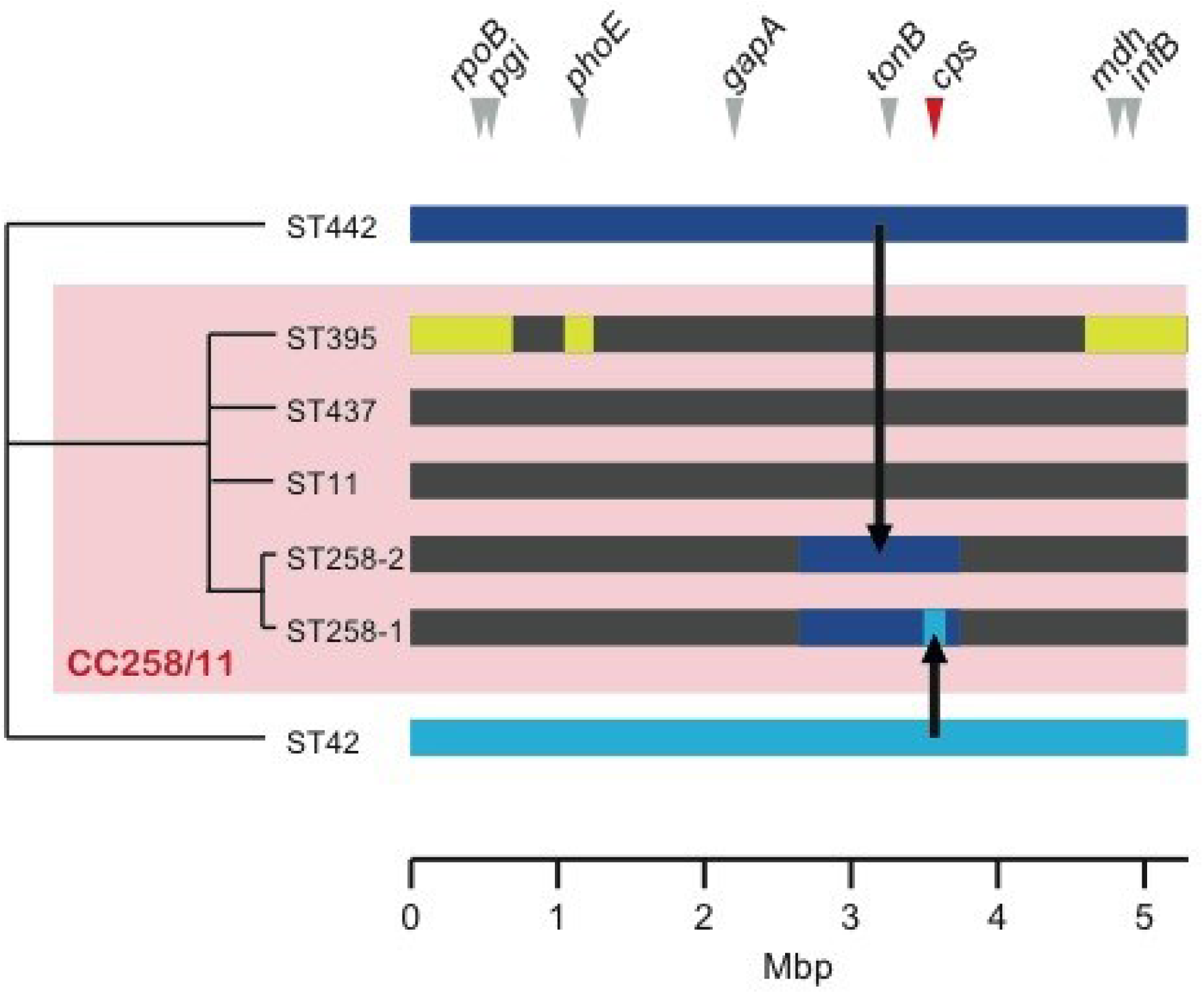
Evolutionary history of the CC258/11 genomes. Dendrogram represents hierarchical relationships among ST442, ST42 and the CC258/11 STs (branch lengths not meaningful). Coloured bars (right) represent the genome of each ST (approximate scale only, coordinates relative to HS11286 genome). Dark grey blocks represent genomic regions descended from the most recent common ancestor of CC258/11; yellow blocks represent recombinant regions acquired from an unknown donor(s); dark blue bar represents the ST442 genome, part of which was imported to ST11 forming the hybrid ST258-2 genome; light blue bar represents the ST42 genome, part of which was imported to ST258-2 forming ST258-1. The positions of MLST loci and the *cps* locus are indicated by arrows.

We identified 11 distinct *cps* loci amongst CC258/11, of which only three had been previously reported in the complex (24), and further IS-associated variants for two of these (**Fig. 4**). The 11 *cps* loci were differentiated by structural (gene content and synteny) rather than nucleotide-level differences. As such, the presence of multiple different variants among otherwise closely related isolates indicates change by horizontal transfer rather than mutation. In contrast to the large recombination events identified in ST395 and ST258, most *cps* locus changes were not associated with large-scale recombination events, suggesting that the recombination breakpoints associated with these capsule switches lay much closer to, or possibly within, the *cps* locus itself. This hypothesis is consistent with the observation of declining sequence conservation towards the centre of the *cps* locus, since this region has likely been affected by a greater number of recombination events than the outer *cps* locus regions throughout the evolutionary history of *K. pneumoniae*. Alternatively, it is possible that the downstream recombination breakpoints lay outside of the *cps* locus but were masked by inter-strain variation at the nearby LPS locus (O antigen locus), which also varied among our *K. pneumoniae* genomes. The latter may be indicative of diversification of LPS within the complex, however as the genetics of LPS production in *K. pneumoniae* are not well understood we are unable to draw conclusions about this from the sequence data.

The capsular serotypes of the ST258 isolates included in this study were unknown, as serotyping is rarely performed for *K. pneumoniae.* However, previous studies have reported expression of serotype K41 among three ST258 *K. pneumoniae* that were tested (39). It is not clear which of the ST258-associated *cps* loci was present in those serotyped isolates, although it was noted that they harboured KPC allele 2, which has been associated with ST258-1/*wzi-29*/cps207-2 (25, 29). The *cps* loci reported here had very few conserved genes (**Fig. 4, 5**) and differed extensively in their complement of sugar-processing genes. Hence while the associated serotypes are not known for all the *cps* loci, exchange of these loci within CC258/11 can be assumed to result in phenotypic capsule switching.

The *cps* loci were correlated with phylogenetically defined sublineages (**Fig. 2**). We also observed diversification of *cps* loci into variants within highly clonal groups, via the activity of IS, which may prove to be useful epidemiological markers for identifying ST258 subclones. The IS involved are diverse and are not site-specific, but were nevertheless observed only in two regions within the *cps* – upstream of *wzi*, or within the central sugar processing region. Only two of the 13 insertions interrupted coding sequences, indicative of selection against transposition events that interrupt expression of the capsular biosynthesis genes (as other IS insertions within CDSs presumably occur but are deleterious in competition with encapsulated strains). However, the IS insertions could hypothetically alter capsular gene expression, or promote capsule switching via rearrangement or horizontal transfer. IS1 and IS5 were identified upstream of *wzi*, within the promoter region of the majority of capsular biosynthesis genes, in four different *cps* loci (**Fig. 4, 5**). It has been shown that insertion of IS5 into specific sites upstream of coding sequence regions can enhance expression of various operons in *E. coli* (40). IS1 insertions in either orientation can also enhance (41) or interrupt (42) gene expression in *E. coli*. Furthermore, insertion of IS1301 upstream of the capsule biosynthesis (*sia*) and export (*ctr*) operons of *Neisseria meningitidis* C has been shown to up-regulate capsule expression and promote resistance to complement-mediated killing (43). Therefore, the apparent hotspot for IS acquisition upstream of *wzi* in the *K. pneumoniae cps* locus may indicate not only purifying selection against deleterious mutants resulting in loss of capsular expression, but also positive selection for enhanced capsule expression at the transcriptional level. In addition, IS1, IS2, IS5 and IS630 were all found within the sugar-processing region of the *cps* loci. IS1-mediated rearrangement of the *cps* locus has been reported in *E. coli* (44); consequently the accumulation of IS in this central region may contribute to capsule switching and diversification of the *cps* loci in *K. pneumoniae*.

While capsular switching within clones is well-understood in Gram-positive pathogens such as *Streptococcus pneumoniae*, extensive capsular variation in Gram-negative *Enterobacteriaceae* has not been widely reported (45). The high number of distinct *cps* locus variants within our sample suggests that capsule switching may be a common event across the wider *K. pneumoniae* CC258/11 (and among other clones such as ST42 for which we identified four genomes with three different *cps* loci: one with *wzi*-29, one with *wzi*-41 and two with *wzi*-33 (all unknown serotypes; genome data in BIGSdb, http://bigsdb.web.pasteur.fr). It is reasonable to assume that as the ST258 sublineages continue to evolve, their *cps* loci will continue to diversify through recombination, transposition and potentially transposase-mediated horizontal gene transfer. The pool of potential *cps* locus donors is expansive; 78 capsular serotypes have been defined, and *wzi* and *wzc* sequencing efforts suggest that many more *cps* loci exist in the wider *K. pneumoniae* population (27, 28). The ST442/*wzi-154* and ST42/*wzi-29 K. pneumoniae* we identified as being closely related to the donors involved in ST258 recombination events, were isolated under very different circumstances to those described in previous reports (ST442/*wzi-154* from bovine infection in the US and ST42/*wzi-29* from human bacteremia in Singapore). This serves as a reminder that circulating genetic variation, including virulence determinants such as the *cps* locus, can be disseminated through both clinical and environmental sources. The latter niche is currently drastically under-sampled, meaning that much of the genetic variation circulating within global *K. pneumoniae* populations is not yet captured.

Given that *K. pneumoniae* capsule variants are immunologically distinct (46), the ability of KPC CC258/11 to undergo frequent capsular switching is concerning, and suggests that capsular-based vaccines may be of limited use. One possible strategy would be to target a wider range of capsule types, similar to the approach currently used in the design of *S. pneumoniae* capsular vaccines (currently 10 or 13 types). However, comprehensive population surveillance would be required in order to monitor the response to vaccination, which in the case of *S. pneumoniae* has included the emergence of vaccine escape strains (47) and the expansion of pre-existing clones expressing capsule types not targeted by the vaccine (48). A more comprehensive understanding, including investigation of diversity and evolution of capsular loci amongst the broader population of clinical, human carriage and environmental *K. pneumoniae* isolates, is required.

## Materials and Methods

### Genome sequence data analysed in this study

Genomic data representing 39 *K. pneumoniae* CC258/11 representatives were included in this study (**Table 1**). Genome assemblies for 30 isolates were retrieved from GenBank. Sequence reads for one isolate (KpMDU1) were generated on the Ion Torrent platform. 76 bp paired-end (PE) sequence reads for eight isolates were generated on the Illumina Genome Analyzer GAII platform as part of a global diversity study of *K. pneumoniae*. Accessions for all CC258/11 sequence data are given in Tables 1 and 2.

**Table 2.**
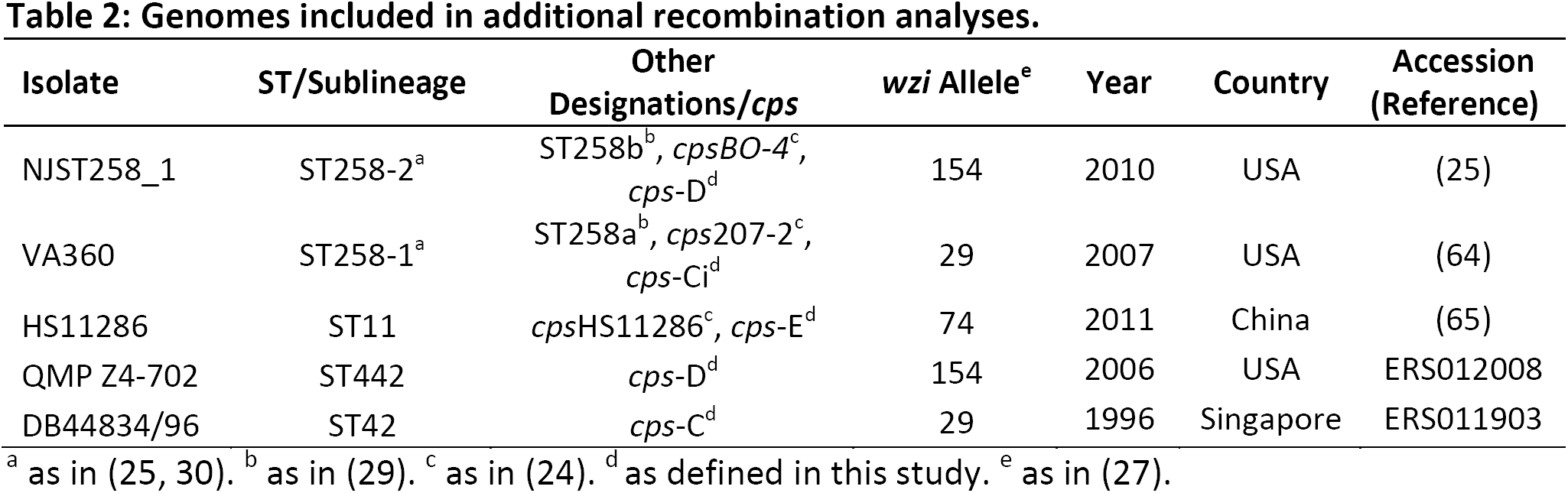
Genomes included in additional recombination analyses.

| Isolate | ST/Sublineage | Other Designations/*cps* | *wzi* Allele<sup>e</sup> | Year | Country | Accession (Reference) |
| --- | --- | --- | --- | --- | --- | --- |
| NJST258_1 | ST258-2<sup>a</sup> | ST258b<sup>b</sup>, *cpsBO-4*<sup>c</sup>, *cps-D*<sup>d</sup> | 154 | 2010 | USA | (25) |
| VA360 | ST258-1<sup>a</sup> | ST258a<sup>b</sup>, *cps207-2*<sup>c</sup>, *cps-Ci*<sup>d</sup> | 29 | 2007 | USA | (64) |
| HS11286 | ST11 | *cpsHS11286*<sup>c</sup>, *cps-E*<sup>d</sup> | 74 | 2011 | China | (65) |
| QMP Z4-702 | ST442 | *cps-D*<sup>d</sup> | 154 | 2006 | USA | ERS012008 |
| DB44834/96 | ST42 | *cps-C*<sup>d</sup> | 29 | 1996 | Singapore | ERS011903 |

<sup>a</sup> as in (25, 30). <sup>b</sup> as in (29). <sup>c</sup> as in (24). <sup>d</sup> as defined in this study. <sup>e</sup> as in (27).

For genome-wide SNP screening of potential CC258/11 members or recombination donors, two public read sets were also analysed (ERP000165, ERP002642).

#### Genome-wide phylogenetic analysis of K. pneumoniae

Sequence reads or, for public genomes, 100 bp PE reads simulated from assembled sequences using SAMTools wgsim (49) with 0% error rate were mapped against the HS11286 reference chromosome (accession: NC_016845.1) using Bowtie 2 (50). Single nucleotide polymorphisms (SNPs) were identified using SAMTools (51) as previously described (52). Briefly, positions in which an unambiguous SNP call was made in any isolate with phred quality ≥30 and read depth ≥5 were identified, and consensus alleles (unambiguous homozygous base call with phred ≥20) were extracted from all isolates and concatenated to generate a SNP alignment. For the global phylogeny, a concatenated alignment of all 272,365 core genome SNPs was generated (defined as genome positions conserved in ≥99% of the genomes). A maximum likelihood (ML) phylogeny was generated using RAxML (53) and a general time reversible (GTR) substitution model with gamma model of rate heterogeneity. The tree representing the highest ML score among five runs, each of 100 bootstrap replicates, is shown in **Fig. 1**.

#### Phylogeny construction and recombination detection within CC258/11

For the CC258/11 analysis, *s*equence reads or simulated reads (as above) were mapped against the HS11286 reference chromosome using BWA (54) and SNPs were identified as above. A pseudo-whole genome alignment was generated by replacing reference bases with SNP alleles identified in each genome. Putative recombinant genome regions were identified from this whole genome alignment using BRATNextGen (55), with 100 x 20 iterations and a reporting threshold of p<0.05. A ML phylogeny was generated using RAxML as above, using an alignment of SNPs at those sites that were conserved in ≥98% of CC258/11 genomes but not identified by BRATNextGen as affected by recombination (total 5,476 sites). The tree representing the highest ML score among 10 runs, each of 1,000 bootstrap replicates, is shown in **Fig. 2**. The tree was rooted using the NTUH-2044 ST23 reference genome (accession: NC_012731.1) as an outgroup.

#### Investigation of large-scale recombination events affecting ST258

Variable genome positions were identified by read mapping to the ST258 reference, NJST258_1 (accession: CP006923.1) and variant calling as described above. The resulting SNPs were analysed using ChromoPainter (34), assuming a uniform recombination map and running 10 iterations, maximising over the recombination scaling constant. **Table 2** lists the genomes included in this analysis.

#### Identification and annotation of cps loci

For publicly available assemblies, genome sequences were retrieved from GenBank (accessions in **Table 1**). For short read data, reads were assembled *de novo* using two alternative approaches, SPAdes ((56), with kmers 21, 33, 55, 63, 71) and Velvet with Velvet Optimiser (57). For each isolate, the assembly yielding the smallest number of contigs was used for the analysis (SPAdes assembly for all except DM23092/04 and 09-370B). *Cps* loci were identified and extracted from the assemblies using a custom Python script, whereby BLAST was used to identify sequence regions with homology to the flanking genes *galF* and *ugd* (nucleotide BLAST followed by protein BLAST if no nucleotide-level matches were found). When *galF* and *ugd* were not found on the same contig, the nucleotide sequences from and including *galF* or *ugd* up to the ends of their respective contigs were extracted. In such cases, contig adjacency was manually confirmed by visual inspection of paired-end read mapping to the *galF* and *ugd* contigs (reads were mapped against extracted *cps* locus sequences using BWA (54), sorted and compressed with SAMTools (51), and viewed in Artemis (58)). In three cases, the *galF* and *ugd* contigs could not be joined (i.e. *cps* loci were split across more than two assembly contigs) and additional contigs were identified by BLAST search of published *cps* locus sequences, and confirmed by read mapping. In three cases (VA360, KpMDU1 and 09-370B) a putative contig join could not be confirmed using the mapping approach. In the case of the public genome, ATCC BAA-2146, no reads were available for mapping. The annotated *cps* sequences for these loci therefore contain contig breaks, the positions of which are shown in **Fig. 4**.

*Cps* loci were clustered into groups representing distinct structures by visual comparison using BLAST and ACT (59). Single representatives of each distinct *cps* locus cluster were annotated using Prokka (60) together with a reference protein set derived from published *cps* loci available in GenBank (accessions: AB371289.1, AB198423.1, AB289646.1, AB289648.1, AB289650.1, AB290716.1, AB371296.1). The resulting annotations, available at https://github.com/kelwyres/Kp-cps-loci.git, were then manually inspected and curated. Nucleotide sequences of conserved CDS were extracted and pairwise similarities were calculated using MEGA5 (61). Transposases were annotated using IS Finder (https://www-is.biotoul.fr/). *Wzi* alleles were assigned by comparison to the international *K. pneumoniae* BIGSdb (at http://bigsdb.web.pasteur.fr) using SRST2 (http://katholt.github.io/srst2/; dx.doi.org/10.1101/006627).

#### Pairwise nucleotide differences and gene distance Jaccard scores

Nucleotide differences were calculated using SNP data generated as described above. SNPs representing putative recombinant genomic regions identified by BRATNextGen were excluded. The total set of genes represented among the CC258/11 genomes was identified by mapping to a pan genome sequence for CC258/11. The latter was obtained by using iterative contig comparison to collate a non-redundant set of distinct contig sequences (<95% sequence identity) present in the set of assemblies, which was annotated using Prokka (60). The presence of each gene in each read set was determined from mapping data, with presence defined as coverage of ≥95% of the length of the gene with mean read depth ≥5. Jaccard distances (*J*) were calculated as *J* = *a/b*, where *a* was the number of genes that were different between two genomes (i.e. present in one but not both) and *b* was the total number of genes present in either genome (i.e. present in one or both genomes).

### Acknowledgements

KEH was supported by fellowship #628930 from the NHMRC of Australia. Computational analysis was supported by a Victorian Life Sciences Computation Initiative (VLSCI) grant #VR0082. SB is a Sir Henry Dale Fellow, jointly funded by the Wellcome Trust and the Royal Society (100087/Z/12/Z). We thank Trinh Dao Tuyet (National Hospital for Tropical Diseases, Vietnam), Tse Hsien Koh (Singapore General Hospital, Singapore), Mark Thomas and Peter Ostrum (Countryside Veterinary Clinic, LLP., USA) for the provision of bacterial isolates. We thank the sequencing teams at the Wellcome Trust Sanger Institute for genome sequencing of eight isolates (funded by Wellcome Trust grant #098051 to Wellcome Trust Sanger Institute) and the Microbiological Diagnostic Unit (MDU, University of Melbourne, Australia) for sequencing of the KpMDU1 isolate.

